# Prediction of metabolites associated with somatic mutations in cancers by using genome-scale metabolic models and mutation data

**DOI:** 10.1101/2023.07.26.550699

**Authors:** GaRyoung Lee, Sang Mi Lee, Sungyoung Lee, Chang Wook Jeong, Hyojin Song, Sang Yup Lee, Hongseok Yun, Youngil Koh, Hyun Uk Kim

**Affiliations:** Department of Chemical and Biomolecular Engineering (BK21 four), Korea Advanced Institute of Science and Technology (KAIST), Daejeon 34141, Republic of Korea; Systems Metabolic Engineering and Systems Healthcare Cross-Generation Collaborative Laboratory, KAIST, Daejeon 34141, Republic of Korea; Department of Genomic Medicine, Seoul National University Hospital, Seoul 03080, Republic of Korea; Department of Urology, Seoul National University College of Medicine, and Seoul National University Hospital, Seoul 03080, Republic of Korea; BioProcess Engineering Research Center and BioInformatics Research Center, KAIST, Daejeon 34141, Republic of Korea; Department of Internal Medicine, Seoul National University Hospital, Seoul 03080, Republic of Korea

**Keywords:** Cancer, Oncometabolite, Genome-scale metabolic model, Mutation data, RNA-seq

## Abstract

**Background:** Oncometabolites, often generated as a result of a gene mutation, show pro-oncogenic function when abnormally accumulated in cancer cells. Identification of such mutation-associated metabolites will facilitate developing treatment strategies for cancers, but is challenging due to a large number of metabolites in a cell and the presence of multiple genes associated with cancer development.

**Results:** Here we report the development of a computational workflow that predicts metabolite-gene-pathway sets (MGPs). MGPs present metabolites and metabolic pathways significantly associated with specific somatic mutations in cancers. The computational workflow uses both cancer patient-specific genome-scale metabolic models (GEMs) and mutation data to generate MGPs. A GEM is a computational model that predicts reaction fluxes at a genome scale, and can be constructed in a cell-specific manner by using omics data (e.g., RNA-seq). The computational workflow is first validated by comparing the resulting metabolite-gene (MG) pairs with multi-omics data (i.e., mutation data, RNA-seq data, and metabolome data) from 17 acute myeloid leukemia samples and 21 renal cell carcinoma samples collected in this study. The computational workflow is further validated by evaluating the MGPs predicted for 18 cancer types, by using RNA-seq data publicly available, in comparison with the reported studies. Therapeutic potential of the resulting MGPs is also discussed.

**Conclusions:** Validation of the MGP-predicting computational workflow indicates that a decent number of metabolites and metabolic pathways appear to be significantly associated with specific somatic mutations. The computational workflow and the resulting MGPs will help identify novel oncometabolites, and also suggest cancer treatment strategies.

## Background

Metabolic reprogramming is one of the important hallmarks of cancer, and plays a crucial role in cancer progression and development [1]. A wide range of metabolic shifts occur in cancer cells as a result of environmental changes and mutations introduced to metabolic genes, oncogenes and tumor suppressor genes. Metabolic phenotypes observed from cancer cells include, but are not limited to, aerobic glycolysis where glycolysis is upregulated and lactate is produced in the presence of oxygen (normoxia) [2], increased glutamine metabolism (glutaminolysis) [3], increased mitochondrial biogenesis and activities [4], dysfunctions in mitochondrial metabolism [4], and increased proton production [5].

Identification of oncometabolites has introduced a new paradigm for cancer metabolism studies. Oncometabolites are defined to be metabolites that show pro-oncogenic function when abnormally accumulated in cancer cells [6]. Currently, three different metabolites are conceived as oncometabolites, namely fumarate, succinate, and 2-hydroxyglutarate (both L and D forms) across different cancer types. These oncometabolites can be generated by both endogenous (e.g., genetic mutations) and exogenous factors (e.g., hypoxic condition). Fumarate and succinate are generated by loss-of-function mutations in fumarate hydratase and succinate dehydrogenase, respectively, while D-2-hydroxyglutarate is generated by gain-of-function mutations in isocitrate dehydrogenase. These oncometabolites in common inhibit α-ketoglutarate-dependent dioxygenases, which causes epigenetic dysregulation via hypermethylation of DNA and histone. Various mechanisms by which oncometabolites contribute to tumorigenesis still continue to be characterized.

Various metabolic phenotypes of cancers as a result of gene mutations suggest the possibility of the presence of additional oncometabolites, or metabolites significantly associated with specific somatic mutations. Identification of additional oncometabolites may lead to the development of various treatment strategies for cancers, including diagnostic and/or prognostic biomarkers and anticancer drugs [7]. For anticancer drugs, ivosidenib and enasidenib were developed on the basis of oncometabolites, which inhibit mutated *IDH1* [8] and mutated *IDH2* [9], respectively, in acute myeloid leukemia (AML), thereby suppressing the biosynthesis of D-2-hydroxyglutarate. Identifying additional oncometabolites is now expected to be better addressed with the help of the increasingly available volume of cancer-derived omics data, for example RNA sequencing (RNA-seq) data and the use of computational modeling approaches that can fully utilize omics data for counterintuitive insights. Genome-scale metabolic models (GEMs) can be considered for this challenge, which can simulate a cell-specific metabolism under varied environmental and genetic conditions [10, 11]. A GEM is a stoichiometric computational model that contains comprehensive information on metabolic gene-protein-reaction (GPR) associations in a specific cell, and can be simulated to predict reaction fluxes at a genome scale by using numerical optimization techniques and omics data (e.g., RNA-seq). GEMs have so far been reconstructed for more than 6,000 organisms for both medical and industrial biotech applications [10]. Cancer GEMs have been developed to predict drug targets [12, 13], metabolic angiogenic targets [14], oncometabolites [15], and to analyze the metabolic network of multiple cancers [16, 17].

In this study, we develop a computational workflow to predict mutation-associated metabolites for 25 cancer types by reconstructing 1,056 patient-specific GEMs using the corresponding RNA-seq data released by the Pan-Cancer Analysis of Whole Genomes (PCAWG) Consortium of the International Cancer Genome Consortium (ICGC) and The Cancer Genome Atlas (TCGA) (https://dcc.icgc.org/pcawg) [18] (Fig. 1). The computational workflow developed in this study involves the simulation of cancer patient-specific GEMs that predicts so-called metabolite-gene-pathway sets (MGPs) across the multiple cancer types. MGPs indicate metabolites and metabolic pathways that appear to be significantly associated with specific somatic gene mutations. The computational workflow and the resulting MGPs will lay the groundwork for further extended studies on oncometabolites and cancer treatment strategies.

**Fig. 1.**
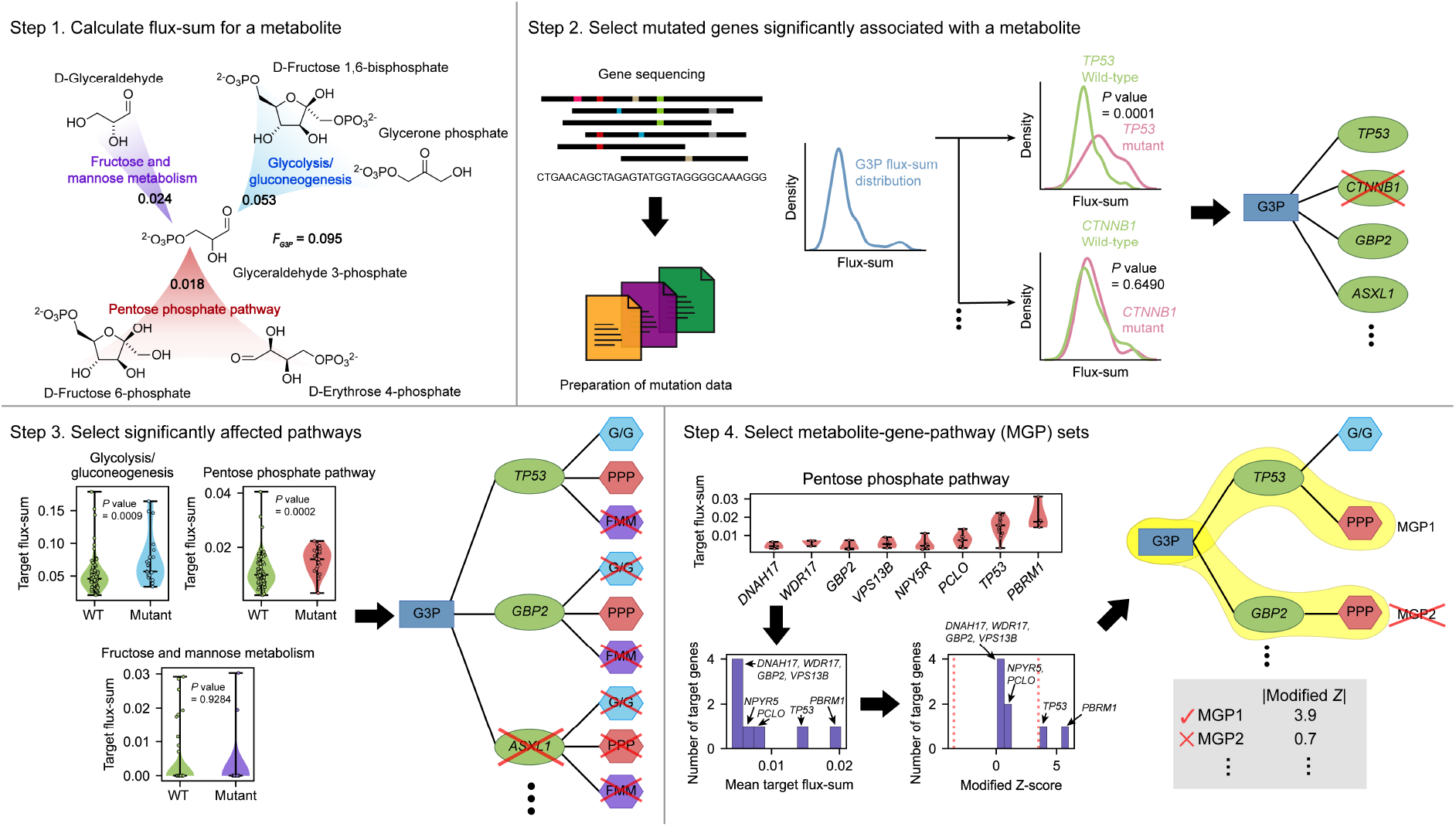
Computational workflow for the prediction of metabolite-gene-pathway sets (MGPs). Computational workflow for predicting MGPs using cancer patient-specific genome-scale metabolic models (GEMs). This workflow is repeated for each metabolite against a list of preselected genes that show mutations in cancers. The computational workflow requires RNA-seq data and mutation data for each cancer sample. Flux-sum value for a target metabolite is first predicted using a cancer patient-specific GEM that is generated using RNA-seq data (step 1). Next, a metabolite is paired with a gene if flux-sum distributions of the metabolite appear to be significantly different upon mutation of the gene (step 2). Metabolite-gene (MG) pairs predicted from the previous step are connected with metabolic pathways that biosynthesize a target metabolite if these pathways show significantly different ‘target flux-sum’ values upon mutation of a target gene (step 3). MG pairs from the previous step are removed if such target pathways are not found. Finally, MGPs are selected by identifying target genes in each target pathway that show target flux-sum values significantly different from those of other target genes in the same pathway (step 4). For this, for each target gene in a target pathway, the mean of its target flux-sum values is calculated, and converted to the modified Z-score. The selected MGPs should have their modified Z-score satisfying the threshold of ‘3.5’.

## Results

### Reconstruction of 1,056 cancer patient-specific GEMs across 25 cancer types

To develop the computational workflow for the prediction of MGPs across multiple cancer types, cancer patient-specific GEMs were first reconstructed using the PCAWG and TCGA RNA-seq data. A previously developed generic human GEM ‘Recon 2M.2’ [12] was integrated with the PCAWG and TCGA RNA-seq data, which attempted to reconstruct GEMs for 1,056 cancer patients that represent 25 cancer types (Fig. 2a). Here, in this study, samples from CNS-GBM and CNS-Oligo were combined to obtain a greater number of *IDH1* mutants from both gliomas that have been relatively well studied for the *IDH1* mutation-associated oncometabolites [19, 20]. The reconstructed GEMs for 7 Eso-AdenoCA and 6 TCGA-LAML samples were discarded in this study because they did not complete up to 24 out of 182 metabolic tasks (Methods), whereas patient-specific GEMs from other cancer types successfully completed all the metabolic tasks. All the reconstructed GEMs were further evaluated using MEMOTE (metabolic model tests) [21], which, as a result, showed a high level of consistency: average scores of 95% for ‘Mass Balance’, 93% for ‘Charge Balance’, and 98% for ‘Metabolite Connectivity’. The resulting 1,043 patient-specific GEMs across the 24 cancer types contained, on average, information on 71.9 metabolic pathways, 3,828.9 reactions, and 1,213.8 unique metabolites (Fig. 2b). Liver-HCC GEMs appeared to have the greatest average number of reactions (i.e., 3964.0 reactions on average), while TCGA-LAML GEMs showed the smallest average number of reactions (i.e., 3448.0 reactions on average); this difference in the model size is likely attributed to unique metabolic activities associated with each cancer type (Fig. 2b). To further confirm that the cancer type-specific GEMs reflect different tissues of origin, reaction contents of the 1,043 reconstructed GEMs were subjected to t-distributed stochastic neighbor embedding (t-SNE) [22], which clearly showed that the GEMs from the same cancer type tend to be better clustered than those from different cancer types (Fig. 2c). This result partly demonstrates the biological quality of the patient-specific GEMs reconstructed in this study.

**Fig. 2.**
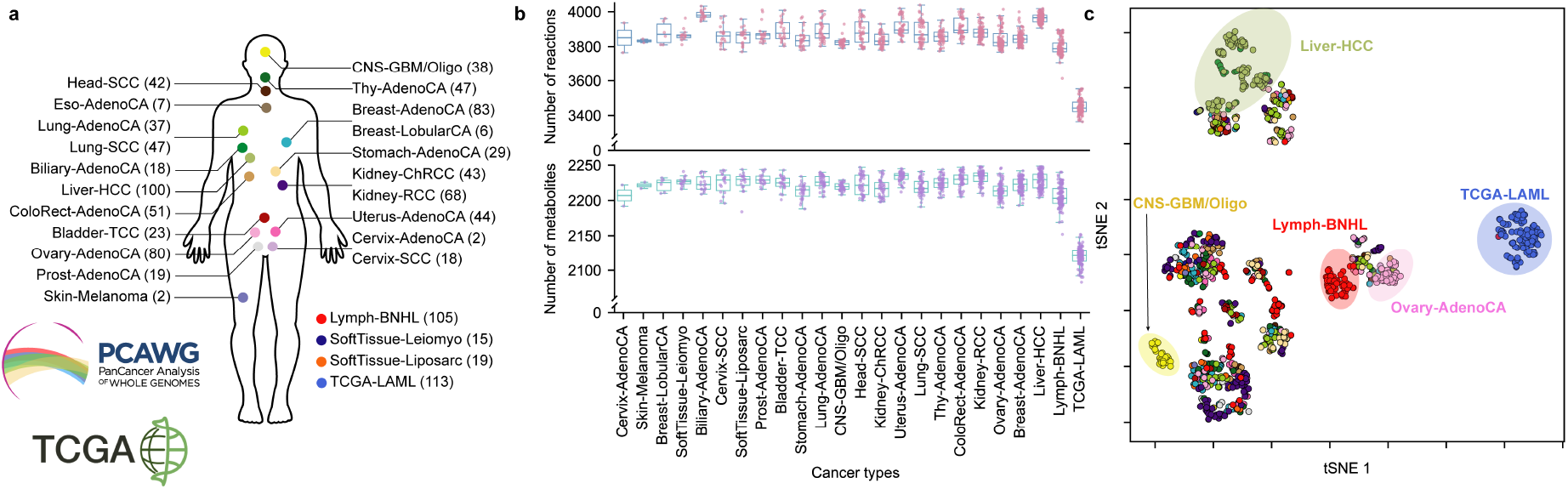
Overview of reconstructing cancer patient-specific genome-scale metabolic models (GEMs). **a** Reconstruction of 1,056 patient-specific GEMs for 25 cancer types by using the PCAWG and TCGA RNA-seq data. The number of samples for each cancer type is presented in a parenthesis next to the cancer type abbreviations. **b** Number of reactions (pink) and metabolites (purple) across the 1,043 GEMs. **c** t-SNE plot of the reaction contents of the 1,043 cancer patient-specific GEMs. Same colors are used as presented in **a**.

### Computational workflow for predicting metabolite-gene-pathway sets (MGPs)

Using the 1,043 patient-specific GEMs and the mutation data from the PCAWG whole genome sequencing (WGS) data and TCGA whole exome sequencing (WES) data for 24 cancer types, MGPs were predicted using a computational workflow that consists of four steps (Fig. 1). This workflow is applied to a metabolite, and generates MGPs as an output. Therefore, this workflow is repeated for entire metabolites of each patient-specific GEM across the 24 cancer types, except for currency metabolites (e.g., ATP and H+). This workflow begins with the calculation of so-called flux-sum value [23] of each metabolite (step 1 in Fig. 1). Flux-sum of a metabolite is defined to be the total sum of all the fluxes necessary for the generation or consumption of that metabolite, which quantifies the intracellular importance of that metabolite. Flux-sum approach was used to examine the robustness of bacterial metabolism [23], predict antibacterial targets [24, 25] and redesign bacterial metabolism for the enhanced chemical production [26].

In the second step, a metabolite was paired with a gene if flux-sum distributions of the metabolite appeared to be significantly different upon mutation of the gene (step 2 in Fig. 1). For convenience, a metabolite and a mutated gene involved in MGP candidates are referred to as ‘target metabolite’ and ‘target gene’ hereafter, respectively. For this metabolite-gene (MG) pairing, PCAWG and TCGA mutation data were prepared, which covered a total of 930 samples, each having 0-586 mutated genes and representing 18 cancer types (Methods). At this stage, as a result, a unique list of 31,521 MG pairs was generated across the 18 cancer types. In this unique list, a compartment for a metabolite was not considered; for example, two pairs, *IDH1* mutant with akg_c and akg_m (α-ketoglutarate in cytoplasm and mitochondria, respectively), were counted as one.

Next, MG pairs predicted from the previous step were connected with metabolic pathways that biosynthesize a target metabolite if these pathways show significantly different ‘target flux-sum’ values upon mutation of a target gene (step 3 in Fig. 1). Here, the target flux-sum value refers to the summation of all the fluxes from a metabolic pathway that contributes to the biosynthesis of a target metabolite. Also, a contributing metabolic pathway considered in MGP candidates is referred to as a ‘target pathway’. MG pairs from the previous step were removed if target pathways were not found. Information from this step was thought to help understand the mechanism behind the association between a target metabolite and a target gene. Indeed, among the MG pairs predicted from the second step, there were pairs that showed a statistical significance (in terms of *P* value) between a target metabolite and a target gene at a genome-scale level, but with no such significance at a pathway level. For example, 2-oxoglutarate was predicted to be significantly affected by *COL6A3* mutation in CNS-GBM/Oligo at a genome scale (in the step 2), but no such significance was observed between 2-oxoglutarate and *COL6A3* mutation at individual 2-oxoglutarate biosynthetic pathways, including: alanine and aspartate metabolism; citric acid cycle; glutamate metabolism; glycine, serine, alanine and threonine metabolism; urea cycle; transport reactions; and additional unassigned reactions. Therefore, ‘2-oxoglutarate-*COL6A3*’ pair was not selected for an MGP from this workflow. Additionally, MG pairs associated with exchange/demand reactions, transport reactions (except for those associated with essential amino acids) or unassigned reactions were not considered because they provide limited information on explaining the biological link between a target metabolite and a target gene; in GEMs, transport reactions are usually annotated with genes at a lower confidence than typical metabolic genes. As a result, 17,656 MGP candidates were generated from this step.

Finally, MGPs were selected by identifying target genes in each target pathway that show corresponding target flux-sum values significantly different from target flux-sum values of other target genes in the same pathway (step 4 in Fig. 1). For this, for each target gene in a target pathway, the mean of its target flux-sum values was calculated, and converted to the modified Z-score (Methods). The resulting modified Z-scores would subsequently reveal target genes that show atypical target flux-sum values despite being in the same target pathway for MGP candidates. For example, 42 MGP candidates involving 42 target genes, all predicted to be associated with 5,10-methenyltetrahydrofolate in Lymph-BNHL, were collected for folate metabolism. Despite their involvement in folate metabolism, only two target genes, *BTK* and *EP300*, encoding Bruton’s tyrosine kinase and histone acetyltransferase p300, respectively, appeared to have the mean flux-sum values significantly different from the other 40 target genes according to the modified Z-scores. Therefore, *BTK* and *EP300* were selected to be final target genes for the target metabolite ‘5,10-methenyltetrahydrofolate’ and folate metabolism in Lymph-BNHL. If fewer than three MGP candidates are available for a target pathway, all the MGP candidates are considered to be significant. From this step, 4,335 MGPs were generated as final sets for the 18 cancer types.

### Evaluation of the MGP-predicting computational workflow using AML and renal cell carcinoma samples

The computational workflow predicting MGPs was first evaluated using multi-omics data from the 17 AML and 21 renal cell carcinoma (RCC) samples. Here, the multi-omics data include mutation data (from either targeted gene sequences or WES data), transcriptome (RNA-seq) and metabolome data; they were experimentally obtained in this study for the AML samples (Methods) and the RCC samples (Methods). This evaluation was in particular focused on whether the computational workflow would generate biologically meaningful MG pairs included in the final MGPs predicted from the 17 AML samples and the 21 RCC samples (Fig. 1). As with the PCAWG data, 17 AML patient-specific GEMs and 21 RCC patient-specific GEMs were first reconstructed using the corresponding RNA-seq data. It should be noted that: one AML patient-specific GEM was discarded in this study because it did not satisfy all the metabolic tasks (i.e., the incapacity to use L-lysine in mitochondria); and among the 21 RCC samples initially collected, RNA-seq data was not properly generated for one RCC sample due to the too low RNA sample purity, and therefore 20 RCC patient-specific GEMs were generated as a result. Subsequently, the reconstructed 16 AML GEMs and 20 RCC GEMs were subjected to the computational workflow (Fig. 1). In this evaluation, seven mutated somatic genes were preselected and considered for the 16 AML samples (Fig. 3a), and another six mutated somatic genes were considered for the 20 RCC samples (Fig. 3e).

**Fig. 3.**
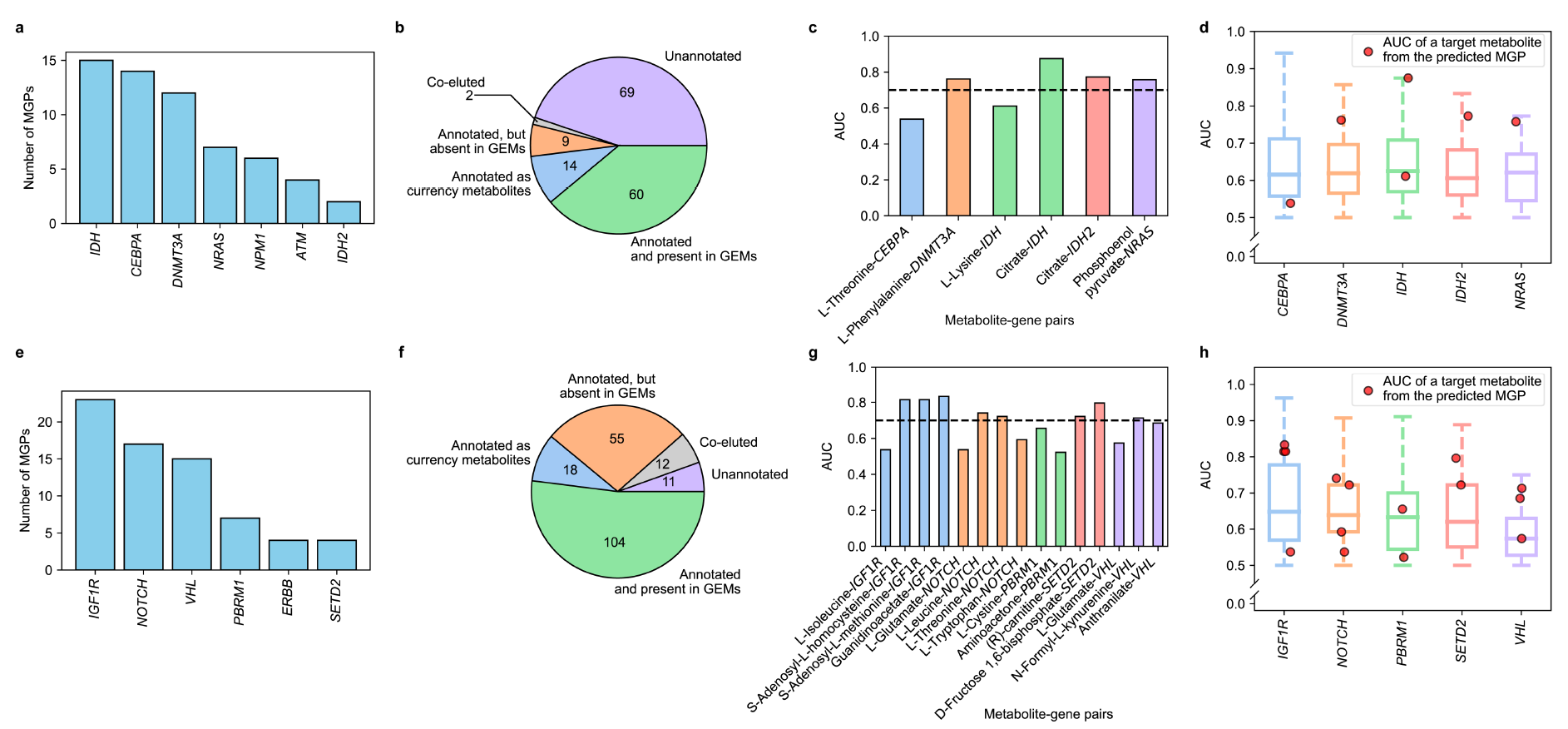
Analysis of metabolite-gene (MG) pairs from metabolite-gene-pathway sets (MGPs) predicted for the 16 AML samples and 20 RCC samples. **a** Number of MGPs predicted for the seven mutated genes from the 16 AML samples. It should be noted that samples having *IDH1* or *IDH2* mutation were also considered together, presented as ‘*IDH’*, in order to examine the overall effects of mutations in *IDH1* and *IDH2*. **b** Classification of the detected peaks from relative quantification of metabolites from the 17 AML samples. **c** AUC values of target metabolites from the final six MGPs, which were predicted from the computational workflow and supported with the AML metabolome data (Fig. 1). AUC values of target metabolites were predicted using MetaboAnalyst [28]. The black dashed line indicates the AUC value of 0.7. **d** AUC values for target metabolites from the final six MGPs (red dots) and 60 metabolites from the AML metabolome data; these 60 metabolites include those not predicted as a target metabolite for MGPs, and are paired with each of the presented target genes (box plots). These 60 metabolites correspond to the peaks in the metabolome data that are annotated, and also present in the GEMs in (**b**). **e-h** Same analyses (a-d) conducted for the 21 RCC samples. In **e**, MGPs were predicted for the six mutated genes from the 20 RCC samples. Samples having *NOTCH1* or *NOTCH2* mutation were considered together as ‘*NOTCH*’, and samples having *ERBB2*, *ERBB3* or *ERBB4* mutation were considered together as ‘*ERBB*’ in order to collect the sufficient number of samples to generate AUC values. In **h**, AUC values for target metabolites from the final 15 MGPs (red dots) and 104 metabolites from the RCC metabolome data are presented. These 104 metabolites include those not predicted as a target metabolite for MGPs, and are paired with each of the presented target genes (box plots).

With 355 unique metabolites with flux-sum values from the 16 AML GEMs, 60 MGPs involving 59 MG pairs were predicted from the computational workflow (Fig. 3a). Five target metabolites (i.e., citrate, L-lysine, L-phenylalanine, phosphoenolpyruvate and L-threonine) that belong to six MG pairs out of the final 59 MG pairs were detected in the 17 AML metabolome data (Fig. 3b,c). Next, biological significance of the target metabolites detected in the AML metabolome data was examined whether these target metabolites would show significantly different concentrations, depending on mutation of a target gene across the AML samples. The significance of a metabolite is presented in terms of the area under the receiver operating characteristic (ROC) curve (AUC), a metric often used as a discriminating power for biomarkers [27], by using MetaboAnalyst [28] (Methods). Among the final six MG pairs supported with the metabolome data, target metabolites paired with *DNMT3A*, *IDH*, *IDH2* or *NRAS* showed AUC values greater than 0.7 [29, 30] (Fig. 3c). Here, it should be noted that the samples having the *IDH1* or *IDH2* mutation were also considered together, presented as ‘*IDH*’, in order to examine the overall effects of mutations in both *IDH1* and *IDH2*. AUC values of the target metabolites in these four MG pairs also appeared to be mostly higher than AUC values of 60 metabolites detected in the metabolome data (Fig. 3d). These results revealed that the computational workflow played a role in selecting biologically more meaningful MG pairs in the final MGPs.

Similar conclusion was also derived from evaluation of the computational workflow using the RCC samples. By applying the computational workflow to the 20 RCC GEMs, 70 MGPs including 69 MG pairs were initially predicted (Fig. 3e); 14 target metabolites involved in 15 out of the final 69 MG pairs were detected in the 21 RCC metabolome data, which allowed the same evaluation as the MG pairs from the AML samples (Fig. 3f,g). As a result, eight out of the 15 MG pairs showed AUC values greater than 0.7 (Fig. 3g). Also, the target metabolites in these eight MG pairs mostly showed greater AUC values than 104 metabolites detected in the metabolome data (Fig. 3h).

### MGPs predicted for the 18 cancer types

A total of 4,335 MGPs from the computational workflow across the 18 cancer types were next evaluated. Overall, Lymph-BNHL generated the greatest number of MGPs (534 MGPs), followed by Liver-HCC (368 MGPs), Breast-AdenoCA (364 MGPs), and Lung-SCC (356 MGPs) (Fig. 4a). These cancer types also had the greatest number of mutated genes among the 18 cancer types except for Breast-AdenoCA: 244, 231, and 221 mutated genes for Lymph-BNHL, Liver-HCC, and Lung-SCC, respectively. There were also cancer types that had a relatively high number of mutated genes despite a small number of samples (e.g., Lung-SCC and LungAdenoCA in Fig. 4a), and the opposite (i.e., greater number of samples than mutated genes; e.g., Ovary-AdenoCA, Kidney-RCC, CNS-GBM/Oligo and TCGA-LAML in Fig. 4a) was also observed. Regarding the number of MGPs predicted, Lymph-BNHL showed a substantially greater number than Liver-HCC although these two cancer types had similar numbers of samples and mutated genes. Moreover, Breast-AdenoCA showed a similar number of MGPs as Liver-HCC although Liver-HCC had almost twice the number of mutated genes than Breast-AdenoCA. This statistics suggests that the resulting MGPs were not necessarily biased by the number of samples and mutated genes. Interestingly, oncogenes and tumor suppressor genes appeared to be slightly more associated with the MGPs than other target genes across the 18 cancer types.

**Fig. 4.**
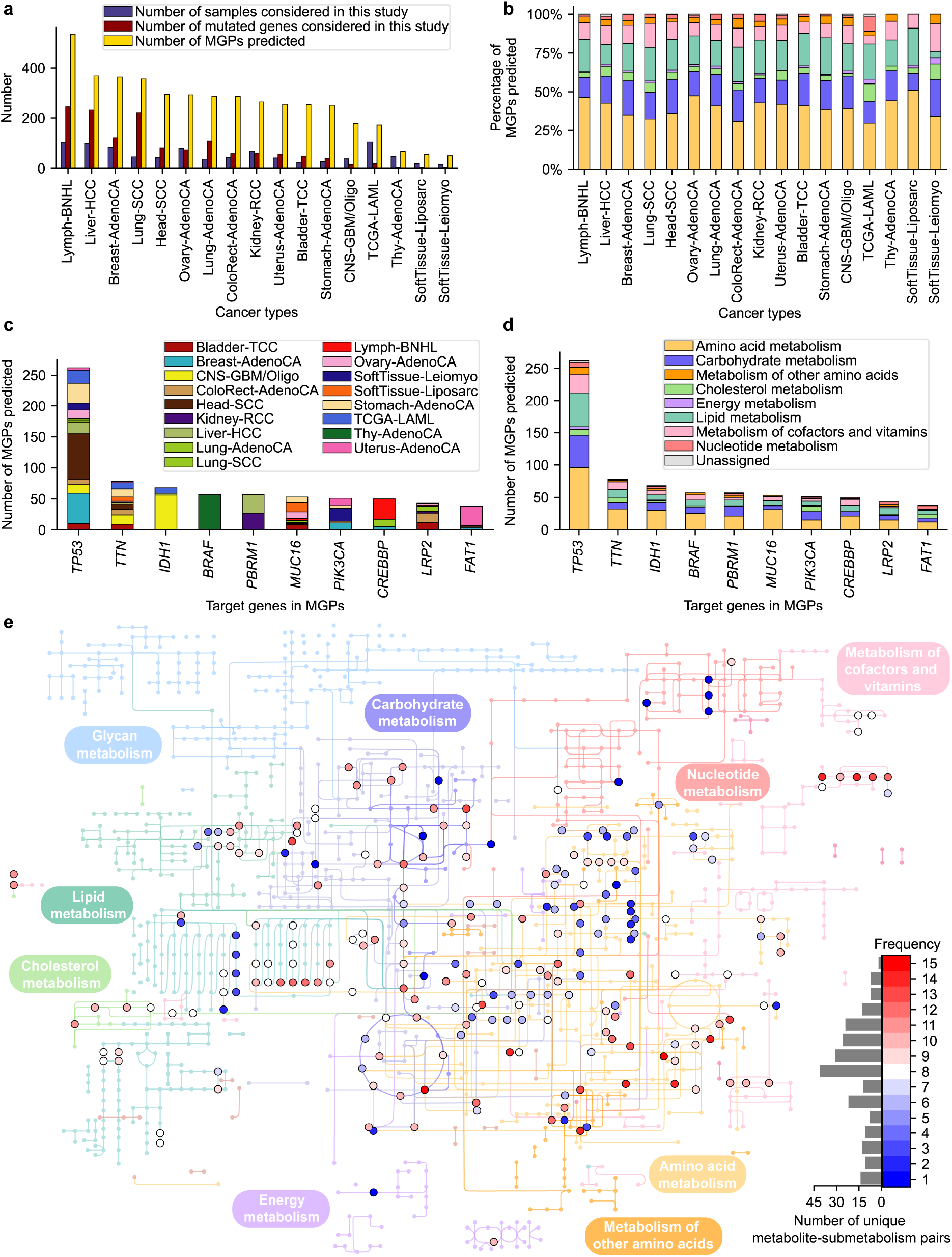
Overview of the predicted MGPs across 18 cancer types. **a** Number of samples and mutated genes considered in this study, and the number of predicted MGPs for each cancer type. **b** Percentages of submetabolisms (on the basis of KEGG pathways) where MGPs were predicted for each cancer type. Colors in bar graphs indicate submetabolisms that are presented in (**d**). **c and d** Ten target genes associated with the greatest number of MGPs where (**c**) the number of cancer types and (**d**) the number of submetabolisms are presented for each target gene. **e**, Distribution of target metabolites associated with the MGPs predicted for the 18 cancer types across the genome-scale human metabolic pathways. Target metabolites and submetabolisms related to target pathways in the MGPs are presented in the metabolic map without target genes. Frequency, shown with different colors between blue and red, refers to the number of cancer types where a target metabolite appeared.

Next, the predicted 4,335 MGPs were categorized into eight different submetabolisms according to the target pathways to gain better insights into these MGPs. As a result, in each cancer type, MGPs were mostly shown to belong to amino acid metabolism (38.5% of MGPs on average for the 18 cancer types), followed by carbohydrate metabolism (19.1%) and lipid metabolism (18.9%) (Fig. 4b). The results are overall consistent with the knowledge of cancer metabolism: for example, increased intracellular concentration of L-leucine associated with *KRAS* mutation in amino acid metabolism [31], and generation of D-2-hydroxyglutarate (carbohydrate metabolism) [19] and altered cholesterol homeostasis (lipid metabolism) [32] as a result of the *IDH1* mutation. Interestingly, the percentage of the predicted MGPs associated with lipid metabolism was remarkably different between two sarcomas, SoftTissue-Liposarc and SoftTissue-Leiomyo, and this different metabolic composition appeared to be consistent with their biology [33]; SoftTissue-Leiomyo (leiomyosarcoma) occurs in smooth muscle [34], whereas SoftTissue-Liposarc (liposarcoma) appears in adipocytes [35]. Cell growth of the liposarcoma is highly affected by fatty acid biosynthesis, which has been suggested as a therapeutic target [36].

A closer look into the target genes involved in the predicted MGPs across the 18 cancer types further showed that seven out of the top ten target genes were cancer driver genes: *TP53*, *IDH1*, *BRAF*, *PBRM1*, *PIK3CA*, *CREBBP*, and *FAT1* [37] (Fig. 4c). The MGPs associated with these target genes appeared to be involved in multiple cancer types with the exception of *BRAF*-associated MGPs. *BRAF*-associated MGPs were predicted to occur solely in Thy-AdenoCA, and also, *IDH1*-associated MGPs mostly appeared in CNS-GBM/Oligo. As expected, these driver genes all appeared to be associated with multiple submetabolisms through MGPs with the three most representative submetabolisms being amino acid metabolism, carbohydrate metabolism and lipid metabolism (Fig. 4d). Representative target metabolites from these three submetabolisms (Fig. 4e) were: 4-aminobutanal and 2-oxoglutarate (predicted in 15 out of 18 cancer types), and 4-aminobutanoate, L-lysine, putrescine, (3*R*,5*S*)-1-pyrroline-3-hydroxy-5-carboxylate, and *trans*-4-hydroxy-L-proline (14 out of 18 cancer types) from amino acid metabolism; D-fructose 6-phosphate and 6-phospho-D-gluconate (13 out of 18 cancer types), and acetyl-CoA, glyceraldehyde 3-phosphate, and 2-oxoglutarate (12 out of 18 cancer types) from carbohydrate metabolism; and decanoyl-CoA, dodecanoyl-CoA, and octanoyl-CoA (12 out of 18 cancer types) from lipid metabolism. Taken together, MGPs predicted from the 18 cancer types overall appeared to be in good agreement with the existing knowledge of cancer metabolism.

It has been known that the same gene mutation can show different metabolic effects in different cancer types [38]. To examine this idea, the MGPs predicted to be associated with *PBRM1*, *PIK3CA*, *CREBBP* or *FAT1* were further examined. Indeed, the different metabolic effects of the same gene mutation were observed, depending on a cancer type, for these four target genes. For example, *PBRM1*-associated MGPs predicted for Kidney-RCC and Liver-HCC showed that histidine metabolism and fatty acid biosynthesis in Kidney-RCC appeared to be affected by *PBRM1* mutation in contrast to pentose phosphate pathway for Liver-HCC. Some of these predicted MGPs were supported by previously reported experimental evidences, including deregulation of histidine metabolism in Kidney-RCC with *PBRM1* mutation [39], and decreased availability of cholesterol upon *PIK3CA* mutation in human breast epithelial line (MCF10A) [40]. Thus, it is expected that the MGPs predicted herein can serve as a reference for further examining the different metabolic effects of gene mutations that have not been experimentally validated.

The novel MGPs predicted across the multiple cancer types may also have a therapeutic potential as supported by following examples. First, a MGP ‘L-leucine-*BRCA1*-transport, extracellular’ was predicted for Ovarian-AdenoCA. L-Leucine activates mTOR pathway [41], which has been suggested as a therapeutic target for *BRCA1*-deficient cancer [42]. Thus, L-leucine restriction in the diet may help treat ovarian cancer with *BRCA1* mutation by less activating mTOR pathway. Next, two MGPs, ‘phosphoenolpyruvate-*PIK3CA*-glycolysis/gluconeogenesis’ and ‘fumarate-*PIK3CA*-citric acid cycle’, were predicted for Breast-AdenoCA, and may provide hypotheses for overcoming trastuzumab resistance in breast cancer with *PIK3CA* mutation [43]. One study showed that trastuzumab resistance might be treated by targeting altered glucose metabolism [44], and, in accordance with the two MGPs, both phosphoenolpyruvate and fumarate were reported to be more available in trastuzumab-resistant gastric cancer [45]. Thus, controlling the availability of phosphoenolpyruvate and/or fumarate may contribute to treat trastuzumab-resistant breast cancer with *PIK3CA* mutation. Another three MGPs paired with *VHL*, a cancer driver gene frequently mutated in RCC [46], support the inhibition of indoleamine 2,3-dioxygenase 1 (IDO1) as a drug target, which was previously attempted [47]. IDO1 converts L-tryptophan to N-formyl-L-kynurenine in tryptophan metabolism, and the three predicted MGPs are: ‘N-formyl-L-kynurenine-*VHL*-tryptophan metabolism’ and ‘anthranilate-*VHL*-tryptophan metabolism’ for the RCC samples collected in this study, and ‘N-formylanthranilate-*VHL*-tryptophan metabolism’ for Kidney-RCC. IDO1 inhibition can stabilize tryptophan metabolism that is often upregulated in RCC, and causes immunosuppression [46]. Finally, two MGPs, ‘reduced glutathione-*KEAP1*-glutamate metabolism’ from Lung-AdenoCA and ‘L-leucine-*KRAS*-transport, extracellular’ from ColoRect-AdenoCA, are well aligned with previous drug target suggestions: inhibition of glutaminase in lung adenocarcinoma with *KEAP1* mutation [48], and inhibition of *LAT1* (or *SLC7A5*) encoding ‘solute carrier family 7 member 5’ in colorectal cancer with *KRAS* mutation [49], respectively. These evidences suggest that the predicted MGPs are not only consistent with the knowledge of cancer metabolism, but also provide reasonable treatment strategies, especially drug targets.

### MGPs predicted for CNS-GBM/Oligo

Finally, MGPs predicted for CNS-GBM/Oligo were further analyzed in comparison with the reported studies on these cancers. This comparative analysis would reveal specific MGPs that agree with previous findings as well as novel MGPs that can be validated in future. First, generation of D-2-hydroxyglutarate as a result of the *IDH1* mutation has been well studied in gliomas [19]. Indeed, this finding was well captured by the MGPs predicted for the CNS-GBM/Oligo (Fig. 5a), which included ‘akg-*IDH1*-citric acid cycle’ and ‘akg-*IDH1*-glutamate metabolism’ from the computational workflow. It should be noted that ‘akg’, a direct precursor of D-2-hydroxyglutarate, was paired with *IDH1* because D-2-hydroxyglutarate is not reflected in the generic human GEM. In glioblastoma cells having the *IDH1* mutant, pyruvate [50], glutamate [50], lactate [51], and choline [52] were also found to show different intracellular concentrations, or biosynthetic reactions for these metabolites were shown to have different activities, compared to the counterpart cells having the wild-type *IDH1* (Fig. 5a). These previous findings except for citrate were all consistent with the MGPs predicted for CNS-GBM/Oligo.

**Fig. 5.**
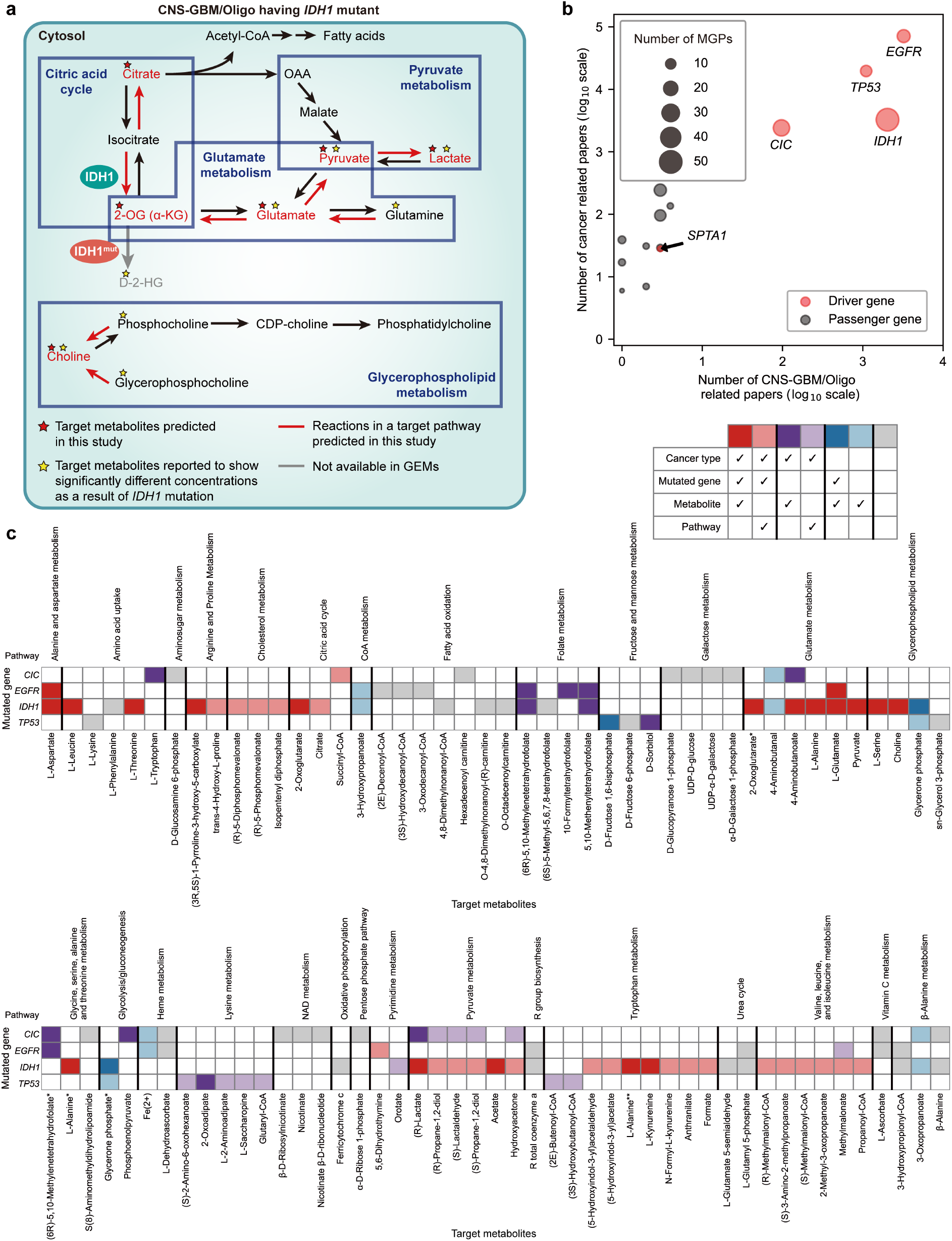
MGPs predicted for CNS-GBM/Oligo. **a** *IDH1*-associated target metabolites (red stars) in the MGPs predicted for CNS-GBM/Oligo, and those with experimental evidence from previous studies (yellow stars). This map shows the overall consistency between the *IDH1*-associated target metabolites predicted and the reported studies. Red lines indicate reactions in a target pathway involved in the predicted MGPs. Black and grey lines indicate reactions available and those unavailable in the generic human GEM Recon 2M.2, respectively. **b** Number of the retrieved papers on 13 target genes in the MGPs predicted for CNS-GBM/Oligo and various cancers in general. Pink and grey circles indicate target genes known to be cancer driver genes and passenger genes, respectively. The size of each circle indicates the number of the MGPs predicted for CNS-GBM/Oligo. **c** Distribution of previous relevant studies for each MGP predicted for CNS-GBM/Oligo. Different colors of the cell represent different levels of consistency between the predicted MGPs and the reported studies, as defined in the table in the upper right-hand corner. White cells indicate that no MGPs were predicted for that particular combination of a gene, a metabolite and a pathway. If a metabolite belongs to two or more pathways, an asterisk (*) is added at the end of its name below the heatmap; if a metabolite belongs to two metabolic pathways, that metabolite with the second appearance is labeled with a single asterisk, and two asterisks for that metabolite with the third appearance.

Next, to understand the volume of previous studies on target genes associated with the MGPs predicted for CNS-GBM/Oligo, papers on 13 target genes from the predicted MGPs were retrieved from PubMed (as of May 2021; Fig. 5b). This paper retrieval was implemented twice, once with an additional keyword of ‘glioma’ and the second round with ‘cancer’. The paper retrieval showed that the previous studies on gliomas appeared to be largely focused on four target genes (*CIC*, *EGFR*, *IDH1* and *TP53*), all cancer driver genes [37], among the 13 target genes (Fig. 5b). This overall pattern was also observed in papers on various cancers in general. Thus, further in-depth analysis of the MGPs was conducted with focus on these four target genes.

For the predicted 115 MGPs that are associated with mutation of at least one of these four cancer driver genes, 79 MGPs (69%) were supported by previous studies to a varying degree with respect to a cancer type and the three MGP components, namely mutated gene, metabolite and pathway (Fig. 5c). Among the four cancer driver genes, MGPs predicted for *IDH1* mutant showed the highest literature coverage (80.4%; 45 out of the 56 predicted MGPs supported by previous studies), followed by *TP53* mutant (78.6%; 11 out of the 14 MGPs supported), *CIC* mutant (48.1%; 13 out of the 27 MGPs supported) and *EGFR* mutant (55.6%; 10 out of the 18 MGPs supported). The highest coverage of the MGPs involving *IDH1* was expected because of this enzyme’s involvement in the generation of D-2-hydroxyglutarate that has been relatively well-studied. The *IDH1* MGPs were predicted to largely affect amino acid-related pathways, in particular glutamate metabolism, which is consistent with the previous studies on this enzyme in cancer cells (Fig. 5c). High coverage of the *TP53* MGPs also recapitulates the metabolic regulation exerted by this gene; the affected target pathways include fructose and mannose metabolism as well as lysine metabolism (Fig. 5c). In contrast to *IDH1* and *TP53*, *CIC* and *EGFR*, encoding capicua transcriptional repressor and epidermal growth factor receptor, respectively, showed relatively lower coverage for their MGPs predicted (grey cells in Fig. 5c), which suggests future research opportunities. Despite the small number of supporting papers, several MGPs predicted for *CIC* and *EGFR* seem to be reasonable in consideration of the biological role of these target genes. For example, three MGPs were predicted to affect fatty acid oxidation upon mutation of the *EGFR* gene, which can be easily inferred from a previous finding that *EGFR* is known to regulate lipid biosynthesis in glioblastoma [53].

*IDH* genes are also frequently mutated in AML, and hence, their mutation affects AML metabolism [54]. To this end, *IDH*-associated MGPs predicted for CNS-GBM/Oligo and TCGA-LAML were compared to examine the metabolic effects of *IDH* mutation in these two cancer types. As expected, α-ketoglutarate, a precursor of D-2-hydroxyglutarate, was predicted as a target metabolite in the MGPs from both CNS-GBM/Oligo and TCGA-AML. However, overall, the predicted MGP profiles were very different in these two cancer types. Metabolites involved in pentose phosphate pathway were observed only in the MGPs from TCGA-AML, and all the other target metabolites were specifically observed in the MGPs from CNS-GBM/Oligo, including those in tryptophan metabolism. A total of seven MGPs associated with *IDH1* and tryptophan metabolism were predicted for CNS-GBM/Oligo; these predictions are consistent with the reported increased level of kynurenine in tryptophan metabolism [55].

### Prediction of MGPs by using another generic human GEM Human1

Finally, the MGP-predicting computational workflow was evaluated by using another recently released generic human GEM, called Human1 [17], to understand whether the use of a different human GEM would affect the MGPs to be predicted. This evaluation was conducted for the same multi-omics data from the AML and RCC samples that were examined using Recon 2M.2 (Fig. 3). Accordingly, 17 AML patient-specific GEMs and 20 RCC patient-specific GEMs were first reconstructed by using Human1 as a template model. Next, MGPs for the AML and RCC samples were predicted using the computational workflow; for the AML samples, 202 MGPs including 198 MG pairs were predicted (Fig. 6a), and for the RCC samples, 156 MGPs including 143 MG pairs were predicted (Fig. 6e). Seven out of the 198 MG pairs for the AML samples (Fig. 6b) and 14 out of the 143 MG pairs for the RCC samples (Fig. 6f) were supported with the corresponding metabolome data; five target metabolites from the AML samples (Fig. 6c) and eight target metabolites from the RCC samples (Fig. 6g) showed AUC values greater than 0.7. Overall, AUC values of these target metabolites appeared to be substantially greater than those of other metabolites from the AML and RCC metabolome data (61 and 111 metabolites, respectively; Fig. 6d,h). As a conclusion, implementation of the computational workflow using Human1 also generated biologically meaningful MG pairs in the final MGPs. However, as expected, use of the two different generic GEMs generated different profiles of the MGPs; for example, for the AML samples, the use of Recon 2M.2 predicted more MGPs with *IDH* than Human1 (Fig. 3a and Fig. 6a). Greater similarities were observed between Recon 2M.2 and Human1 for the RCC samples as the target metabolites with AUC values greater than 0.7 were commonly associated with *IGF1R*, *NOTCH*, *PBRM1*, *SETD2* or *VHL* (Fig. 3h and Fig. 6h).

**Fig. 6.**
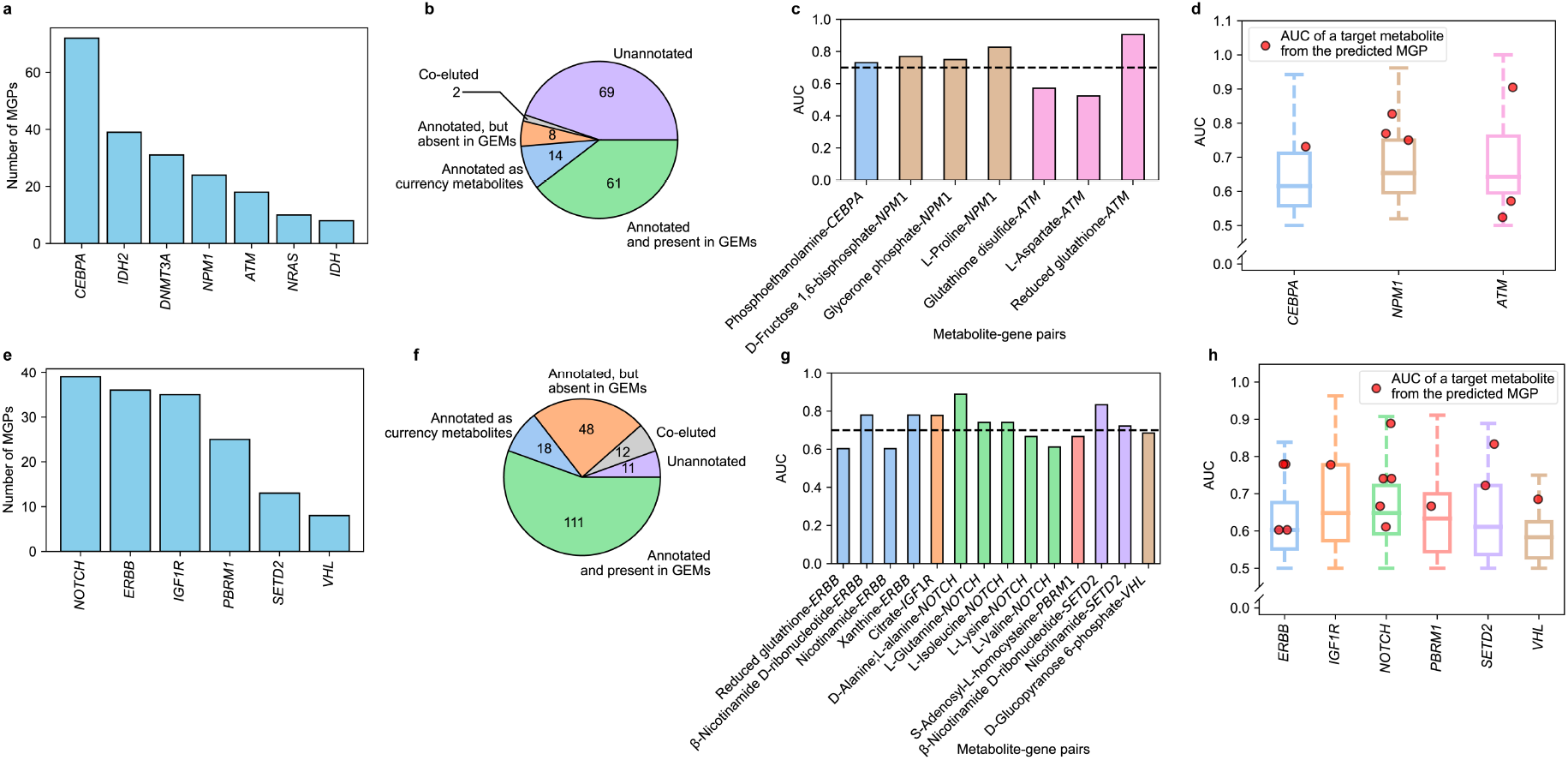
Analysis of MG pairs from MGPs predicted using Human1 for the 16 AML samples and the 20 RCC samples. The same analysis was performed by using Human1 as a template model for the data presented in Fig. 3. **a** Number of MGPs predicted for the seven mutated genes from the 16 AML samples. As in Fig. 3a, the samples having *IDH1* or *IDH2* mutation were also considered together, presented as ‘*IDH*’. **b** Classification of the detected peaks from relative quantification of metabolites from the 17 AML samples. **c** AUC values of target metabolites from the final seven MGPs. The black dashed line indicates the AUC value of 0.7. **d** AUC values for target metabolites from the final seven MGPs (red dots) and 61 metabolites from the AML metabolome data; these 61 metabolites include those not predicted as a target metabolite for MGPs, and are paired with each of the presented target genes (box plots). These 61 metabolites correspond to the peaks in the metabolome data that are annotated, and also present in the GEMs in (**b**). **e-h** Same types of data presented in (a-d) for the 21 RCC samples. In **e**, MGPs were predicted for the six mutated genes from the 20 RCC samples. As in Fig. 3e, the samples having *NOTCH1* or *NOTCH2* mutation were considered together as ‘*NOTCH*’, and the samples having *ERBB2*, *ERBB3* or *ERBB4* mutation were considered together as ‘*ERBB*’. In **h**, AUC values for target metabolites from the final 14 MGPs (red dots) and 111 metabolites from the RCC metabolome data are presented. These 111 metabolites include those not predicted as a target metabolite for MGPs, and are paired with each of the presented target genes (box plots).

## Discussion

In this study, we investigated the possible presence of metabolites that are significantly associated with specific somatic mutations in multiple cancer types by using GEMs and mutation data. For this, RNA-seq data from PCAWG and TCGA representing 25 different cancer types as well as the AML and RCC samples were first used to reconstruct cancer patient-specific GEMs. Subsequently, the computational workflow involving the GEMs and the mutation data of the cancer patients was developed that generates so-called MGPs that present metabolites and contributing metabolic pathways that are significantly associated with somatic mutations in cancers. This computational workflow was first validated by using the multi-omics data (i.e., mutation data, RNA-seq data, and metabolome data) from the 17 AML and 21 RCC samples; the same analysis was also conducted for breast cancer samples by using their multi-omics data previously reported [56]. The MGPs predicted for 18 cancer types were analyzed in regard to their metabolic effects and therapeutic potential. Furthermore, the MGPs predicted for CNS-GBM/Oligo were extensively compared with findings from the reported studies. This validation process showed that the computational workflow developed in this study generates reliable MGPs, which can serve as candidate targets for further in-depth studies. Finally, the computational workflow was also demonstrated by using another generic human GEM Human1, which again generated biologically meaningful MGPs.

Despite our efforts, the computational workflow developed in this study can be further updated by addressing several challenges. First is to collect a greater number of samples, preferably for more diverse cancer types, which would allow more rigorous validation of the computational workflow. A major challenge here is to collect a balanced number of cohorts, each having specific gene mutations of interest, to obtain various metabolites associated with each of these mutations. Each cohort will obviously have a highly varied mutational landscape, which would involve the unforeseen effects of complex gene-gene interactions and mutation types on metabolite profiles. Another challenge is to generate multi-omics data (e.g., mutation data, RNA-seq data and metabolome data) for a greater number of samples from various cancer types. This will allow more rigorous validation of the computational workflow, and systematic analysis of cancer type-specific metabolism. For these reasons, this computational workflow is not intended for an immediate clinical application, for example detecting a cancer biomarker in a person. Rather, it is hoped that the computational workflow and its resulting MGPs serve as the groundwork for identifying novel oncometabolites, and for facilitating the development of various treatment strategies.

## Conclusions

In this study, we developed the computational workflow that uses GEMs and mutation data of the cancer patients in order to predict metabolites and metabolic pathways that are significantly associated with specific somatic mutations in cancers. By using RNA-seq data from PCAWG and TCGA, 4,335 MGPs were predicted for the 18 cancer types. First, the computational workflow was validated by using the multi-omics data (i.e., mutation data, RNA-seq data, and metabolome data) from the 17 AML and 21 RCC samples that were collected in this study. Comparison of the resulting MG pairs with the multi-omics data revealed a decent number of metabolites that showed significant changes in their concentration as a result of specific gene mutations. The MGPs predicted for 18 cancer types were also thoroughly examined in comparison with the reported studies, in particular whether they are overall consistent with the knowledge of cancer metabolism and the therapeutic potential previously suggested. Further rigorous analysis was made on the MGPs predicted for CNS-GBM/Oligo. Overall, the validation studies showed that the predicted MGPs are biologically meaningful, which can serve as candidate targets for further in-depth studies. The computational workflow developed in this study can also be considered for other cancer types not covered in this study upon availability of the relevant datasets (i.e., mutation data and RNA-seq data).

## Methods

### Generation of personal GEMs using RNA-seq data

A previously developed generic human GEM Recon 2M.2 [12] was transformed into a context-specific (personal) GEM through the integration with personal RNA-Seq data from the acute myeloid leukemia (AML) samples, the renal cell carcinoma (RCC) samples, TCGA, or PCAWG. Task-driven Integrative Network Inference for Tissues (tINIT) method, along with a rank-based weight function, was used to generate personal GEMs [11, 12, 57]. A total of 182 metabolic tasks were evaluated for the resulting personal GEMs. All the resulting personal GEMs were evaluated using MEMOTE [21]. Another generic human GEM Human1 [17] was also used to generate the cancer patient-specific GEMs by using the AML and RCC RNA-seq data.

### Visualization of cancer patient-specific GEMs

Metabolic reaction contents of the resulting 1,043 cancer patient-specific GEMs were visualized using t-SNE to cluster the GEMs according to their cancer type. To implement t-SNE, an input binary vector was prepared for each GEM, indicating the presence and absence of a reaction as ‘1’ and ‘0’, respectively. For hyperparameters, ‘number of principal components’ and ‘perplexity’ were set to be ‘30’ and ‘20’, respectively.

### Calculation of flux-sum values of metabolites

Flux-sum values for each metabolite were calculated for each personal GEM reconstructed in this study. For this, intracellular fluxes were first predicted by minimizing the distance between transcript expression level (or gene expression level for TCGA-LAML data) from RNA-seq data and target reaction fluxes to be calculated in an objective function; target reactions in the objective function were determined through transcript-protein-reaction associations (or GPR associations for TCGA-LAML data), and the least absolute deviation method was implemented for this distance minimization as previously described [12]. Next, flux-sum (*F_i_*) of metabolite *i* in each GEM was calculated according to a previously defined mathematical formulation [23]:

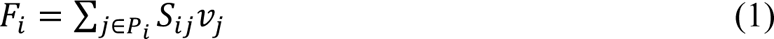

where *S_ij_* refers to the stoichiometric coefficient of metabolite *i* involved in reaction *j* at a reaction rate *v_j_*, and *P_i_* for a set of reactions producing metabolite *i*. Reactions consuming metabolite *i* were not considered when predicting MGPs.

### Preparation of AML and RCC samples

Both bone marrow samples and RCC samples (primary kidney cancer samples) were collected at Seoul National University Hospital. Bone marrow samples were obtained from 17 patients diagnosed with acute myeloid leukemia (AML) from 2016 to 2019. RCC samples were obtained from 21 patients diagnosed with RCC from 2016 to 2021. These samples were subjected to targeted gene sequencing or whole exome sequencing (WES) as described below.

### Targeted sequencing of AML and RCC samples

Mutation data for the AML and RCC samples were partly obtained from the targeted gene sequencing. For the AML samples, mutation data were obtained from SNUH FIRST Hemic Treatment Panel, which is a targeted gene panel consisting of 76 genes that are recurrently mutated in myeloid neoplasms; these 76 genes were sequenced using next-generation sequencing. Fifty ng of DNA collected from bone marrow samples from patients with hematologic malignancy was used for targeted sequencing. Library preparation was performed according to SureSelect^QXT^ Target Enrichment system (Agilent Technologies). Finally, paired-end 150 bp sequencing was conducted using NextSeq 550Dx system (Illumina). For the RCC samples, mutation data were obtained from SNUH FIRST Cancer Panel that covers information on 148 genes. For these samples, 50-200 ng of DNA was collected from the RCC samples, and the same sequencing protocol above was also implemented.

### Whole exome sequencing of AML and RCC samples

Mutation data for the AML and RCC samples were additionally obtained from WES. Four AML samples and 12 RCC samples were subjected to WES. For exome sequencing, 50 Mb targeted exons were captured using SureSelect Human All Exon V5 (Agilent Technologies). Hundred bp paired-end sequence reads of the captured exons were generated using HiSeq 2000 Sequencing System (Illumina) according to the manufacturer’s instructions.

### Bioinformatics analysis of DNA sequences

The WES data, the SNUH FIRST Hemic Treatment Panel data and the SNUH FIRST Cancer Panel data were analyzed using SNUH First Panel Analysis Pipeline. First, the FASTQ files were subjected to quality control, and only those that met the criteria were further analyzed. Pair-end alignment to the human genome reference hg19 was performed using Burrows-Wheeler Alignment (BWA) 0.7.17 [58] and Genome Analysis Toolkit (GATK) Best Practices [59]. After finishing the alignment step, an ‘analysis-ready’ BAM files were generated, and SNV and InDel were detected using GATK UnifiedGenotyper 4.1.9 [59], SNVer 0.5.3 [60] and LoFreq 2.1.2 [61]. Detected variants were annotated using SnpEff 5.0 [62] with RefSeq, COSMIC, dbSNP, ClinVar, and gnomAD as reference databases.

### RNA-seq analysis of AML and RCC samples

Total RNA was isolated from each AML and RCC sample using PAXgene Blood RNA Kit (Qiagen). RNA integrity and concentration for library preparation were determined by using 2100 Bioanalyzer (Agilent Technologies). TruSeq Stranded mRNA (Illumina) was used to prepare RNA-seq libraries. RNA-seq libraries were quantified with KAPA Library Quantification Kit (Kapa Biosystems) according to the manufacturer’s library quantification protocol. The 151 bp paired-end sequencing of these libraries was performed using NovaSeq 6000 Sequencing System (Illumina). FastQC 0.10.1 [63] was used to evaluate the quality of raw reads. RNA-seq reads were aligned to the human genome reference hg19 using Spliced Transcripts Alignment to a Reference (STAR) 2.7.0f [64]. Uniquely aligned reads were counted using featureCounts 1.6.2 [65]. Finally, expression levels of each transcript were estimated in transcripts per million (TPM).

### Metabolome analysis of AML and RCC samples

Metabolome analysis was conducted at Human Metabolome Technologies (HMT) by using capillary electrophoresis time-of-flight mass spectrometry measurement for the relative quantification of metabolites. The AML and RCC samples for metabolome analysis were prepared in accordance with instructions from HMT. For the AML samples, 354 peaks, covering 243 peaks from Cation mode and 111 peaks from Anion mode, were detected, and among them, 185 peaks were annotated on the basis of HMT’s standard library and ‘Known-Unknown’ peak library. For the RCC samples, 363 peaks, covering 243 peaks from Cation mode and 120 peaks from Anion mode, were detected; the 363 peaks were annotated using the same libraries as the AML samples.

The resulting metabolome data were further processed and analyzed using MetaboAnalyst 5.0 (http://www.metaboanalyst.ca) [28]. First, a metabolite was not considered in this study if its corresponding data appeared to be missing in more than 20% of the AML and RCC samples [66]. For the remaining metabolites, their missing values were imputed by using *k*-nearest neighbors. Upon this initial processing, 154 peaks survived from 354 peaks for the 17 AML samples, and 200 peaks survived from 363 peaks for the 21 RCC samples. The relative quantification data for each metabolite were additionally subjected to three types of normalization, including sample normalization via ‘normalization by sum’, data transformation via ‘generalized logarithm’, and data scaling (i.e., autoscaling) via ‘mean centering’ together with ‘division by the standard deviation’. Finally, ‘Classical univariate ROC curve analysis’ was used to generate AUC values for metabolites as a function of a gene mutation in the 17 AML samples and the 21 RCC samples. For the 17 AML samples, a total of 94 peaks were excluded, including 69 unannotated peaks, two co-eluted peaks, nine annotated peaks absent in Recon 2M.2, and 14 peaks annotated as currency metabolites (Fig. 3b). For the 21 RCC samples, a total of 96 peaks were excluded, including 11 unannotated peaks, 12 co-eluted peaks, 55 annotated peaks absent in Recon 2M.2, and 18 peaks annotated as currency metabolites (Fig. 3f).

### Preparation of RNA-seq data from PCAWG and TCGA

A total of 1,056 cancer patient-specific RNA-Seq data and their corresponding mutation data across 25 cancer types were obtained from Pan-Cancer Analysis of Whole Genomes (PCAWG) Consortium of the International Cancer Genome Consortium (ICGC) and The Cancer Genome Atlas (TCGA) [18].

### Preparation of mutation data from PCAWG WGS data and TCGA WES data

For the 943 cancer patient-specific WGS data from PCAWG and 113 cancer patient-specific WES data from TCGA, following genes were discarded in this study: mutations covered by fewer than seven alternative reads in a sample; synonymous mutations; genes having mutations that occur in fewer than three samples in a cancer type; wild-type genes in fewer than three samples in a cancer type; and ‘subset’ gene mutations.

### Processing flux-sum values for predicting MGPs

In the second step of the computational workflow predicting MGPs, flux-sum profiles of cancer patient-specific GEMs were categorized into wild-type and mutant groups for a mutated gene in a cancer type. To obtain flux-sum values that are significantly different between the wild-type and mutant groups, flux-sum values (*F_i_*) were normalized using quantile normalization method [67] for each cancer type. If a normalized flux-sum value 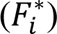 appears to be non-zero for a metabolite despite the original flux-sum value being zero, zero value was used for that metabolite. Flux-sum values of a metabolite between the wild-type and mutant groups were considered significantly different if *P* value from the two-sided Wilcoxon rank-sum test was less than 0.05, which, as a result, allowed pairing a metabolite with a mutated gene for MGP candidates.

In the third step of the computational workflow for selecting ‘target pathways’ that significantly contribute to the biosynthesis of a ‘target metabolite’, ‘target flux-sum values’ were first adjusted in accordance with the normalized flux-sum values of target metabolites in order to preserve the relative ratio of target flux-sum values across contributing pathways that produce a given target metabolite.

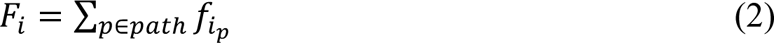

where *f_i_p__* denotes the target flux-sum of pathway *p* producing metabolite *i*, and *path* for a set of pathways producing metabolite *i*. Based on this, target flux-sum of pathway *p* was adjusted as follows:

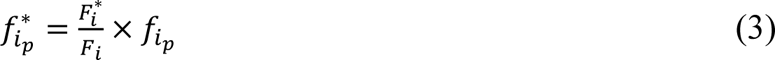

Statistical significance of the target flux-sum values for a target pathway between the wild-type and mutant groups was also examined using the two-sided Wilcoxon rank-sum as in the second step (*P* value < 0.05).

For the final step of the computational workflow, the mean target flux-sum value for each target gene in each target pathway associated with MGP candidates was converted to the modified Z-score:

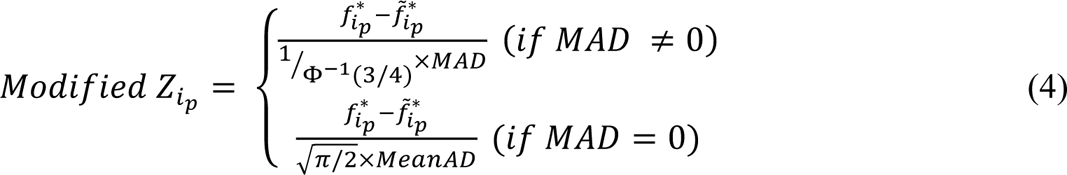

where 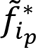 denotes the median adjusted target flux-sum values, Φ denotes the cumulative distribution function of normal distribution, and *MAD* and *MeanAD* stand for median absolute deviation from the median and mean absolute deviation from the median, respectively. A threshold of |modified Z*_i_p__*-score| > 3.5 was considered for a target gene in a MGP candidate to be significant [68].

### Computing environment

Reconstruction and simulation of all the personal GEMs were conducted in Python environment with Gurobi Optimizer 9.0.2 and GurobiPy package (Gurobi Optimization, Inc.). Reading, writing and manipulation of the COBRA-compliant SBML files were implemented using COBRApy 0.6.0 [69]. All the statistical tests were conducted using SciPy 1.4.1 [70]. Principal component analysis initialization and t-SNE were conducted using *scikit-learn* 0.20.3 [71]. Paper retrieval from PubMed was conducted using Biopython 1.74 [72]. All the plots presented in this study were generated using seaborn 0.10.0 [73] and matplotlib 3.2.0 [74]. The metabolic pathway map in Fig. 4e was generated using Cytoscape 3.8.1 [75] on the basis of human metabolic pathway maps from KEGG [76].

## Declarations

### Ethics approval and consent to participate

This study was conducted according to the Declaration of Helsinki, and was approved by the institutional review board of Seoul National University Hospital (IRB No. H-2107-203-1239 for the AML samples, and H-2203-119-1308 for the RCC samples) and KAIST (IRB No. KH2021-158). All patients gave informed consent at the time of sample collection.

### Consent for publication

Not applicable.

## Availability of data and materials

RNA-seq data from the AML samples and the RCC samples are available at Sequence Read Archive (accession number: PRJNA757576). The PCAWG dataset and the TCGA dataset are available at the ICGC data portal (https://dcc.icgc.org/releases/PCAWG) and the GDC data portal (https://portal.gdc.cancer.gov/), respectively. COBRA-compliant SBML files of 943 cancer patient-specific GEMs for 24 cancer types from PCAWG, 113 cancer patient-specific GEMs for TCGA-LAML, 16 AML patient-specific GEMs, and 20 RCC patient-specific GEMs are available at https://doi.org/10.5281/zenodo.7296304. Source code for the computational workflow predicting MGPs (Fig. 1) is available at https://bitbucket.org/kaistsystemsbiology/mgp_prediction.

## Authors’ contributions

S.Y.L., H.Y., Y.K. and H.U.K. conceived the project. G.L. and S.M.L. conducted all the computational studies. All the authors analyzed the data. G.L., S.M.L., S.Y.L., H.Y., Y.K. and H.U.K. wrote the manuscript.

## Acknowledgments

We are grateful to Seokhyeon Kim for his assistance with preparing documents for the institutional review board.

